# PTM-Shepherd: analysis and summarization of post-translational and chemical modifications from open search results

**DOI:** 10.1101/2020.07.08.192583

**Authors:** Daniel J. Geiszler, Andy T. Kong, Dmitry M. Avtonomov, Fengchao Yu, Felipe V. Leprevost, Alexey I. Nesvizhski

## Abstract

Open searching has proven to be an effective strategy for identifying both known and unknown modifications in shotgun proteomics experiments. Rather than being limited to a small set of user-specified modifications, open searches identify peptides with any mass shift that may correspond to a single modification or a combination of several modifications. Here we present PTM-Shepherd, a bioinformatics tool that automates characterization of PTM profiles detected in open searches based on attributes such as amino acid localization, fragmentation spectra similarity, retention time shifts, and relative modification rates. PTM-Shepherd can also perform multi-experiment comparisons for studying changes in modification profiles, e.g. in data generated in different laboratories or under different conditions. We demonstrate how PTM-Shepherd improves the analysis of data from formalin-fixed paraffin-embedded samples, detects extreme underalkylation of cysteine in some datasets, discovers an artefactual modification introduced during peptide synthesis, and uncovers site-specific biases in sample preparation artifacts in a multi-center proteomics profiling study.

## BACKGROUND

Database searching of shotgun proteomics data is a commonly used strategy for identification of peptides and proteins from complex protein mixtures (1, 2). Peptide identification in this strategy most commonly relies on matching tandem mass spectrometry (MS/MS)-derived peptide spectra to their theoretical counterparts using MS/MS database search tools, which requires prior knowledge of the potential modifications that might be present in a sample. This is problematic, as proteins can exist in myriad forms outside of their canonical sequences. For example, protein function is commonly modulated by post-translational modifications (PTMs), and additional chemical modifications from sample processing can hinder identification. Because the search space of all potential peptides including their modifications is so large, when using conventional database search strategies, researchers are forced to limit the modifications considered by their searches, leading to large number of unexplained spectra (3- 6).

Open searching, or mass-tolerant searching, is one strategy that allows researchers to expand their search space and reduce the number of unexplained MS/MS spectra. It has proven to be an effective strategy for identifying both known and unknown modifications in shotgun proteomics experiments (3, 4, 7-9). Rather than being limited to user-specified modifications, open searches identify peptides with mass shifts corresponding to potential modifications or sequence variants. These mass shifts do not, however, contain the same information present in closed searches, most importantly the identity of the modification and what amino acids within the peptide sequence may contain it. Deciphering open search results thus requires subsequent computational characterization to recover this information (4, 10-12).

Here we present PTM-Shepherd, an automated tool that calls modifications from open search peptide-spectrum match (PSM) lists and characterizes them based on attributes such as amino acid localization, fragmentation spectra similarity, effect on retention time, and relative modification rates. PTM-Shepherd can also perform multi-experiment comparisons for studying changes in modification profiles under differing conditions. We utilize these profiles in a wide array of situations to show how additional metrics, interexperiment comparisons, and bulk analytical profiles can be helpful in PTM analysis. Overall, we expect that PTM profiles produced by PTM-Shepherd will greatly enhance understanding of the data at both the macro level for quality control and the micro level for specific PTM identification.

## EXPERIMENTAL PROCEDURES

### PTM-Shepherd: mass shift histogram construction

A histogram of identified mass shifts is constructed using all PSMs from the PSM.tsv file (or multiple PSM.tsv files in the case of multi-experiment analysis) generated by the MSFragger/Philosopher pipeline (**Figure 1**). These PSM.tsv files are typically (by default) filtered to 1% PSM-level and 1% protein level FDR using target-decoy counts, as determined by the Philosopher filter command. The widths of each bin the histogram is 0.0002 Da (by default). This histogram is extended by 5 Da on either side of the most extreme values in order to prevent peaks at the maximum and minimum of the histogram from being truncated after smoothing. Random noise between -0.005 Da and 0.005 Da is added to break ties occurring between bin boundaries and mass shifts. After bin assignment, the histogram is smoothed to make peaks more monotonic. Bin weight is distributed across 5 bins (by default), with the weights assigned to each bin being determined by a Gaussian distribution centered at the bin to be smoothed such that 95% of the bin’s weight is distributed between them. Peaks, representing mass shifts of observed modifications, are called from this histogram.

**Figure 1:**
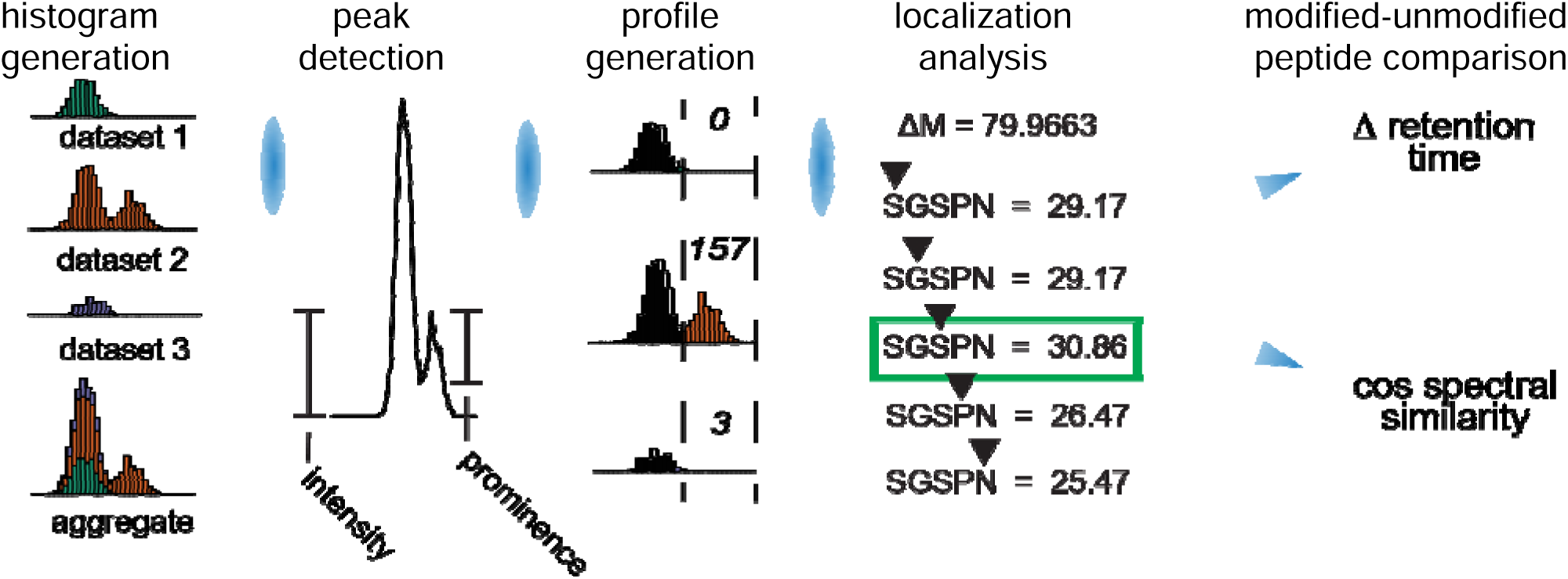
PTM-Shepherd workflow. Data processing begins by aggregating the mass shifts across all datasets into a common histogram. Peaks are determined based on their prominence. The 500 most intense peaks in aggregate are then quantified for each dataset and normalized to size. Peptides with each mass shift are iteratively rescored with the peptide at each position, producing localization scores for each peptide and an aggregate localization enrichment for each mass shift. Finally, modified peptides and their unmodified counterparts are analyzed to have their pairwise cosine spectral similarity and change in retention time calculated.

### PTM-Shepherd: peak picking

PTM-Shepherd picks peaks based on a mixture of peak prominence and signal-to-noise ratio (SNR). Peak SNR is calculated with 0.004 Da sliding window (by default) against a background of 0.005 Da on either side (scaling linearly with peak picking width). A peak’s prominence is calculated from the ratio of its apex to the more intense of either its left or right shoulder, found by following a peak downward monotonically (**Figure 1**). Peaks are called when their prominence exceeds 0.3 (by default). From this list of peaks, the top 500 (by default) are sent to downstream processing. Peak boundaries are considered to be either the observed peak boundary or the defined precursor tolerance, whichever is closer to the apex. PSMs are assigned to the peak if their mass shift falls within the reported peak boundary.

### PTM-Shepherd: mass shift annotation

Detected peaks are iteratively annotated using entries from the Unimod (retrieved: 2 Oct 2019)(13) modification database (including single residue insertions and deletions and isotopic peaks), supplemented with a user-specified list of mass shifts. Each peak is allowed to be decomposed into at most two modifications. Some exceptionally rare or protocol-based modifications (e.g., O18 labeling, N15 labeling) that regularly confounded annotation were removed. Mass differences within 0.01 Da (by default) of a known mass shift are annotated immediately. If a mass shift does not meet this condition, it is then tested against combinations of user-defined mass shifts and known annotations before being checked against combinations of two modifications identified at the previous step above. Failing both of these assignments, mass differences are marked as “Unannotated” and appended to the list of potential modification combinations.

### PTM-Shepherd: mass shift localization

PTM-Shepherd constructs localization profiles for each mass shift peak. Localization profiles are constructed for each experiment, reporting an N-terminal localization rate and a normalized amino acid propensity for each peak. The localization step is performed for every PSM by placing the mass shift at each amino acid in turn and re-scoring the PSM (with the original spectrum) using the same scoring function as in MSFragger. PSMs are considered localizable if there is a position(s) within the peptide sequence that, when the mass shift is placed there, results in more matched fragment ions than using unshifted fragment ions only (i.e. without adding the mass shift anywhere). Localizable PSMs corresponding to the same peak in the mass shift histogram are aggregated, and their characteristics are analyzed. The localization rate for a peak is calculated by counting the number of instances a mass shift was localized to a particular amino acid. If the localization is ambiguous (i.e. several sites scored equally high), the weight of the localization is distributed among all localized residues. Counts are normalized to the rate of localization for a given residue, then divided by each residue’s background content. Background residue content is computed by counting the number of occurrences of each residue in every localizable PSM in the entire dataset (by default). Options for experiment-level normalization at the unique peptide level, and bin-wise normalization at the PSM and unique peptide level are also available.

### Modified-unmodified comparisons

Cosine spectral similarity between modified and unmodified peptides is used to determine how mass shifts affect MS/MS spectra. Unmodified PSMs, i.e. PSMs with a mass shift less than 0.001 Da (by default), are aggregated based on their identified peptide sequence and charge state. If there are more than 50 unmodified spectra for a peptide, 50 are randomly selected for downstream comparisons. Then, for every mass shifted PSM at a given charge state, the average cosine similarity score between this PSM and its corresponding unmodified PSMs at the same charge state is recorded. These similarity scores are aggregated for all PSMs for each mass shift peak, then averaged and reported as that peak’s spectral similarity profile. Retention time effects are also examined. Peptide retention times are extracted from Philosopher’s PSM.tsv output. For every PSM with a mass shift, the average difference in retention time between that PSM and all its corresponding unmodified PSM is calculated. These average retention time differences are aggregated for each mass shift peak, then averaged across all peptides in that peak and reported as that peak’s retention time difference profile.

### Experimental datasets

Four formalin-fixed, paraffin-embedded (FFPE) datasets were used for identifying modifications associated with the fixing process and storage as selected by Tabb et al. for their study (14). Two of these datasets, deemed “Nielsen” (PXD000743) and “Buthelezi” (PXD013107), were acquired on SCIEX TripleTOFs (15). The Nielsen dataset was acquired on a TripleTOF 5600+ and consists of 218449 scans across 20 SCIEX .wiff files, and the Buthelezi dataset was acquired on a TripleTOF 6600 and consists of 474726 scans across 12 SCIEX .wiff files. Two other datasets, deemed “Zimmerman” (PXD001651) and “Nair” (PXD013528), were acquired on Thermo Q-Exactive instruments (16). The Zimmerman dataset consists of 79803 scans across 5 .raw files, and the Nair dataset consists of 245589 scans across 10 .raw files. Files were acquired as .raw or .wiff files and converted to mzML using ProteoWizard’s MSConvert version 3.0.18208.

The synthetic peptide dataset was obtained from ProteomeXchange (PXD004732) in mzML format (17). Only MS runs with the 3xHCD label were included in our analysis. Peptide pools labeled as “SRM” were also excluded. This synthetic peptide dataset consists of unmodified proteotypic human peptides fragmented on a Thermo Fisher Orbitrap Fusion Lumos instrument. Cysteines were incorporated as alkylated cysteines during synthesis.

Additional datasets used in this work were obtained from the Clinical Proteomics Tumor Analysis Consortium (CPTAC) Data Portal in mzML format (18). These were limited to MS runs generated from the CompRef samples, a CPTAC reference material created using breast cancer xenograft pools for quality control and data harmonization purposes. The samples were analyzed using TMT-10 labeling based technology. The first cohort of six experiments (TMT 10-plexes) consists of samples processed at three sites (2 experiments from each site) - the Broad Institute (BI), Johns Hopkins University (JHU), and Pacific Northwest National Laboratory (PNNL) - acquired on an Orbitrap Fusion Lumos as part of the CPTAC harmonization study (19). The second cohort consists of the same CompRef samples processed as longitudinal QC samples as part of three CPTAC datasets: the Clear Cell Renal Cell Carcinoma (CCRCC) dataset generated at JHU (three experiments), the Lung Adenocarcinoma (LUAD) dataset generated at BI (four experiments), and the Uterine Corpus Endometrial Carcinoma (UCEC) dataset generated at PNNL (four experiments). All these data, at all sites, were generated using the CPTAC harmonized data generation protocol (19). These data were processed together using PTM-Shepherd’s multi-experiment setting to generate a single report.

### Database search and statistical validation

All analysis was performed using a database constructed from all human entries in the UniProtKB protein database (retrieved 29 July 2016). Reversed protein sequences were added as decoys and common contaminants from were appended (total targets and decoys: 141,585). Unless specified otherwise, all datasets were processed with the following parameters. Data was searched using MSFragger v2.1(4) with a precursor mass tolerance of +/- 500 Da. Isotope error correction was disabled, and one missed tryptic cleavage was allowed for peptides of 7 to 50 residues in length. Oxidation of methionine was included as a variable modification and cysteine carbamidomethylation was included as a fixed modification. MSFragger mass calibration and parameter optimization was performed for all datasets (20), including fragment ion tolerance. Shifted ions were not used in scoring.

FFPE data was processed using a -200 to 500 Da mass range to match that used in original publication(14). CPTAC data was processed with protein N-terminal acetylation and peptide N-terminal TMT mass of 229.1629 Da as variable modifications (TMT was also specified as fixed modification on Lys). In addition, CPTAC data was searched against combined UniProtKB mouse plus human protein database (retrieved 10 February 2020), with its respective reversed decoys appended to the database, resulting in 252,401 total target and decoy proteins.

PSMs identified using MSFragger were processed using PeptideProphet (21) via the Philosopher v2.0.0 toolkit (22). All processing and filtering was performed on a per-experiment basis. The four FFPE datasets were processed as four experiments. The chloroacetamide-labeled HeLa cells dataset was processed as one experiment containing all 39 fractions. CPTAC samples were grouped experiment-wise, with each experiment containing all 24 fractions. Due to large size, the ProteomeTools synthetic peptide dataset was processed as 11 subsets split based on the five-digit identifier at the beginning of each filename. PeptideProphet parameters for all analyses were default open search parameters: semi-parametric modeling, clevel value set to -2, high accuracy mass mode disabled, masswidth of 1000, and using expectation value for modeling. Resulting PSM matches were filtered to 1% FDR using target-decoy strategy with the help of Philosopher filter command.

## RESULTS AND DISCUSSIONS

### Overview of PTM-Shepherd

The overview of PTM-Shepherd computational workflow is shown in **Figure 1**. The process starts with PTM-Shepherd reading FDR-filtered PSM lists (produced by MSFragger and Philosopher, optionally with open search artifacts removed using Crystal-C (23)), and the mass shift for each PSM, to construct a mass shift histogram (11). After smoothing the histogram, PTM-Shepherd picks peaks based on a mixture of peak prominence and signal-to-noise ratio. From this list of detected peaks, the top 500 (by default) are selected for downstream processing. Basic abundance statistics are then calculated for this list of detected peaks. PSMs are assigned to a particular peak if their mass shift falls within the reported peak boundary, and abundance of the peak is calculated based on spectral counts. PTM-Shepherd can also operate in a multi-experiment mode. In this mode, peak detection is performed on an aggregate mass shift histogram from all experiments, generated from the mass shifts of each experiment weighted according to the proportion of the total PSMs they comprise. The use of a combined histogram for peak detection can greatly simplify comparisons between modifications detected in different conditions and experiments. In this multi-experiment mode, the summary attributes for each detected peak are generated separately for each experiment, and for all data combined.

Once peaks in the mass shift histogram have been called, PTM-Shepherd attempts to determine their identities. Mass shifts are iteratively annotated using entries from the Unimod (13) modification database, isotopic peaks, and user-specified mass shifts, allowing the mass difference to be decomposed into at most two modifications. PTM-Shepherd also constructs localization profiles for each peak. Localization profiles are constructed for each experiment, reporting an N-terminal localization rate and a normalized amino acid propensity for each modification. This analysis is performed for every PSM by placing the mass shift at each amino acid in turn and re-scoring the PSM (with the original spectrum) using the scoring function presented in MSFragger (see Methods).

PTM-Shepherd also computes several metrics that are useful for gaining a better understanding of the nature of those detected mass shifts. For each peak, PSMs containing that mass shift are compared to their unmodified counterparts if present within the same run. First, cosine spectral similarity between modified and unmodified peptides is computed, which is useful for determining how the modifications affect spectra. Then retention time effects are examined, and the average difference in retention time between the peptide with and without modification are reported.

### PTM-palette discovery: analysis of FFPE samples

Understanding how sample processing affects proteins is critical to maximizing their identification. Some techniques – such as formalin-fixing paraffin-embedding (FFPE) – warrant the inclusion of additional modifications to reflect changes in proteins during sample processing (24). Although previous studies have examined which modifications should be included when searching protein data that had been treated with FFPE (24), this was revisited recently by Tabb et al. (14) using a two-pass search. First, an open search was used to identify prevalent mass shifts. Second, they performed a traditional search and informed the localization of their mass shifts with chemical knowledge. We sought to investigate how PTM-Shepherd could be used to validate their findings and streamline this analysis for other datasets and sample preparation protocols.

After their first pass open search, Tabb et al. (14) found five modifications that were consistently present across the four datasets analyzed: mono-methylation, di-methylation, single oxidation, double oxidation, and variable carbamidomethylation. Automated processing with PTM-Shepherd replicates most of these findings. Based on PSM counts and using the same criteria, we find mass shifts of mono-methylation, mono-oxidation, and di-oxidation within the top 10 mass shifts (excluding isotopic peaks) for every dataset (**Supplementary Table 1A**). Interestingly, PTM-Shepherd also finds a notable discrepancy with respect to di-methylation levels. PTM-Shepherd identifies two peaks in close proximity: 27.9954 Da (corresponding to formylation) and 28.0320 (corresponding to di-methylation). Di-methylation is only higher than formylation in one dataset (Nielsen, **Figure 2a**), while the others have formylation between three- and nine-fold higher than di-methylation. To confirm that this was not an artefact of PTM-Shepherd’s signal-to-noise peak picking, we reanalyzed these results with DeltaMass software that implements an alternative (Gaussian mixture modeling) strategy for peak picking (11). For all of these four datasets, DeltaMass found that the region of mass shifts from 27.90 to 28.10 contained two peaks (**Supplemental Figure 1**). For Nielsen, Nair, and Zimmerman, these are easily visible. Even the Buthelezi dataset, while not exhibiting as clear of a separation as the others, places the more abundant peak apex closer to the mass shift value corresponding to formylation. The presence of formylation within a list of most abundant PTMs also makes logical sense given the nature of preservation method.

**Table 1:** Top mass shifts from CPTAC quality control samples. Assigned modifications correspond to automated Unimod matches, with * indicating a manually reannotated mass shift. The “% in Unmodified” column corresponds to the percent of PSMs with a matching unmodified PSM in the unmodified bin. Top two enriched amino acid localizations are shown in columns denoted “AA.”

| Mass Shift | Assigned Modification | PSMs | % in Unmodified | Similarity | Delta RT | N-term rate (%) | AA_1 | AA_2 |
| --- | --- | --- | --- | --- | --- | --- | --- | --- |
| 0.000 | None | 4372080 | 100.00 | 0.98 | 0 | 0 |  |  |
| 1.002 | First isotope | 913405 | 79.43 | 0.81 | 5 | 3 |  |  |
| 229.163 | TMT | 312295 | 74.52 | 0.38 | -70 | 6 | S (5.6) | T (2.4) |
| 2.005 | Second isotope | 177121 | 75.95 | 0.74 | 4 | 1 |  |  |
| -0.984<br>(15.011*) | NH addition to M ** | 156315 | 90.25 | 0.71 | -173 | 1 | M (15.8) |  |
| 230.166 | First isotope + TMT | 141498 | 72.51 | 0.42 | -153 | 6 | S (3.8) |  |
| 0.017 | First isotope +<br>NH addition to M ** | 117922 | 69.97 | 0.78 | 1 | 1 | M (5.8) |  |
| 0.984 | Deamidation | 110967 | 68.84 | 0.68 | 53 | 6 | N (12.7) | R (5.3) |
| 17.026 | Deuterated methyl ester | 92862 | 94.96 | 0.76 | -50 | 1 |  |  |
| 15.011 | Conversion of carboxylic acid to hydroxamic acid | 87013 | 95.22 | 0.54 | -434 | 4 | E (5.5) | D (3.5) |
| -1.002 | Half of a disulfide bridge | 72465 | 81.40 | 0.78 | 16 | 1 | C (4.8) | W (4.5) |
| 15.995 | Oxidation | 59846 | 91.19 | 0.49 | -342 | 21 | W (24.0) | M (11.7) |
| 100.016 | Succinic anhydride labeling reagent | 59070 | 95.76 | 0.55 | 596 | 77 | P (3.6) | S (2.1) |
| 27.995 | Formylation | 52323 | 93.22 | 0.61 | 265 | 61 | S (2.7) |  |
| 79.967 | Phosphorylation | 48379 | 55.30 | 0.64 | 569 | 4 | S (6.2) |  |
| 21.981 | Sodium adduct | 47471 | 98.72 | 0.25 | -148 | 2 |  |  |
| 115.027 | Cleavage product of EGS protein crosslinks by hydroxylamine | 47022 | 97.81 | 0.58 | 610 | 86 | H (2.5) |  |
| 43.010 | Carbamylation | 45450 | 97.10 | 0.64 | 640 | 70 |  |  |
| -18.010 | Dehydration/Pyro-glu from E | 45445 | 97.48 | 0.64 | -175 | 12 | D (3.2) | T (2.5) |
| 1.987 | First isotope + Deamidation | 42988 | 69.08 | 0.66 | 45 | 4 | N (11.9) | R (3.0) |

**Figure 2:**
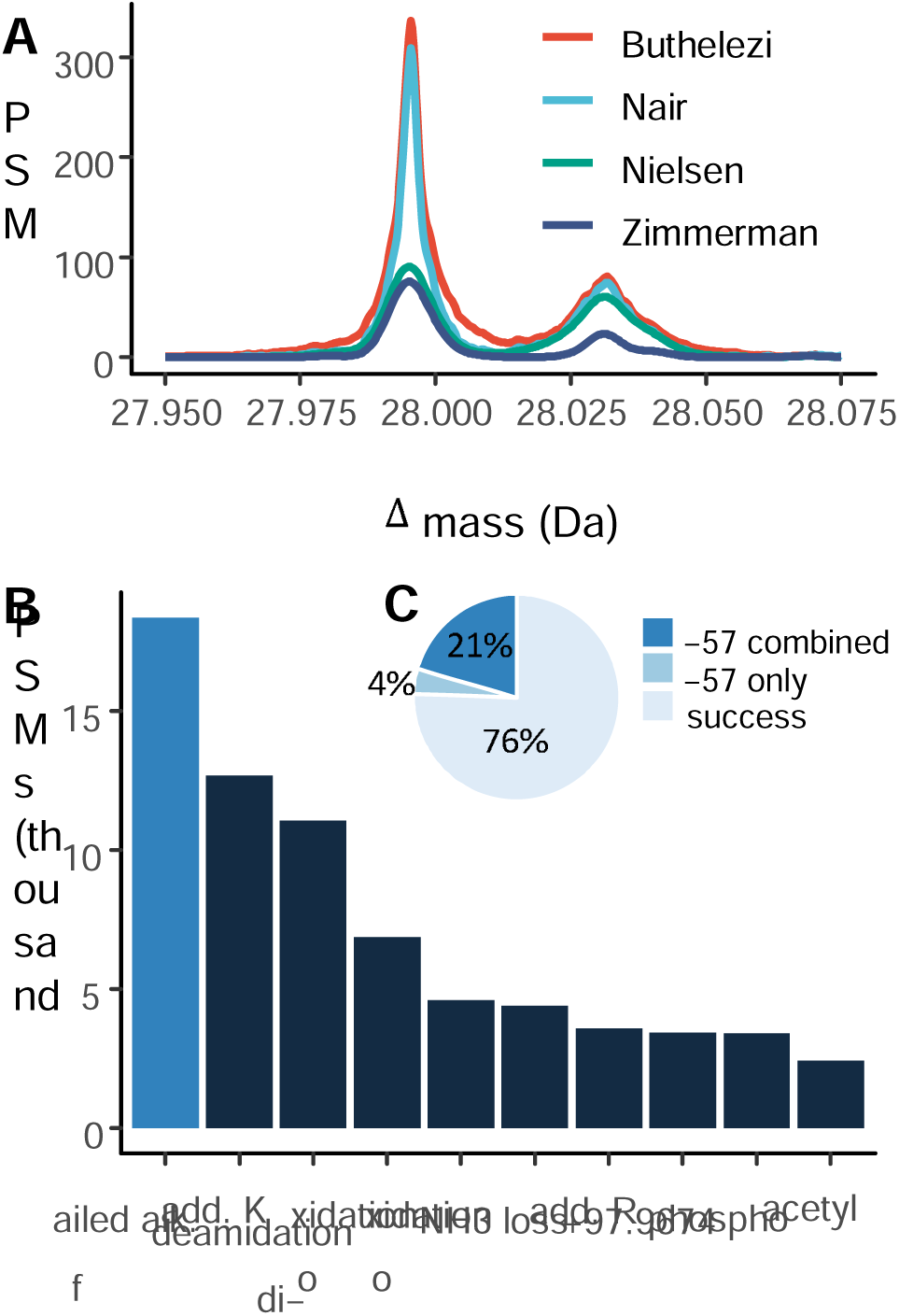
Basic PTM-Shepherd applications. A: PTM-Shepherd identifies two peaks in close proximity for Tabb et al.’s four datasets. All four datasets (Zimmerman, Nair, Nielsen, and Buthelezi) show a mixture of two Gaussian peaks about 28 Da. The consistently more intense peak is at 27.9949, formylation. Only in the Nielsen dataset does di-methylation (28.0313) approach formylation’s intensity. B: PTM-Shepherd identifies more failed alkylation than other common modifications such as deamidation and not-Met oxidation. C: PTM-Shepherd modification decomposition identifies six times as much failed alkylation as is identifiable based on the -57 Da mass shift alone, in total accounting for a quarter of all Cys-containing peptides.

Tabb and colleagues relied on chemical knowledge and other tools (7, 10) to arrive at the final search configuration that included di-oxidation on Met. We chose to investigate this further, using PTM-Shepherd. Because a single oxidation of Met was already included as a variable modification in our open search, a di-oxidation on Met may appear as either a variable modification and a +16 Da mass shift localized to Met or a +32 Da mass shift localized to Met. However, we do not observe enrichment of Met (**Supplementary Table 1A**) localization in either of these instances. In contrast, the enrichment scores for Pro were 9.3 and 5.6 for mono- and di-oxidation, respectively. Outside Pro, we do not see evidence of a clear single source of di-oxidation mass shift. Tabb and colleagues’ gain in the number of PSMs when adding di-oxidation of Met in the search may be explained by the diffuse nature of the modification. When using a closed search strategy with a dynamic +32 Da modification on the Met, any occurrence of two +16 Da events - for example on two alternative oxidation sites - might be interpreted as a Met di-oxidation because peptide ions will converge downstream of the theoretical and experimental oxidation sites. Overall, in our experience in this and other datasets, PTM-Shepherd provides a very reasonable estimate of the most likely modification sites for a particular mass shift.

Formaldehyde adduct (+30.01 Da, annotated as methylol in Unimod) is a known mass shift in FFPE data, and it was detected at high levels in Nair and Zimmerman datasets (in the top 10). According to PTM-Shepherd analysis, this mass shift lacks any significant localization characteristics (localized less than 10% of the time), indicative of a non-covalent adduct. In general, identification of labile modifications is one of the advantages of open database searching with MSFragger compared to other PTM-focused tools or closed searches with variable modifications, all of which are less effective at finding labile modifications that cannot be localized. Summarizing our observations, PTM-Shepherd suggests a slightly modified version of the FFPE PTM palette proposed by Tabb et al.: mono-oxidation of Met and Pro (+15.9949 Da), di-oxidation on Pro (+31.9898 Da), formylation on Lys and N-termini (+27.9949 Da), and monomethylation on Lys and N-termini (+14.0157 Da). It may also be beneficial to include the methylol (+30.01) adduct, but using the mass offset search option of MSFragger that allows both shifted and non-shifted fragment ions in scoring rather than as variable modification (4, 8).

### Detection of Cysteine artefacts following underalkylation

Cysteine is an extremely reactive amino acid, frequently picking up an array of chemical modifications when exposed (25). Unspecified mass shifts, such as those resulting from chemical modifications of Cys, confound peptide identifications and lead to lower recovery rates. Cys alkylation restricts the number of chemical derivatives it can form and prevents interference from disulfide bonds, and as such has been a mainstay of proteomics processing for decades (26). Chloroacetamide (CAA) and iodoacetamide (IAA) are the two most common alkylating reagents used in proteomics workflows. Previous comparisons of these reagents found that IAA generally has a higher rate of cysteine alkylation than CAA when applied at the same concentration, but with the caveat of higher rates of off-target effects (27). Here, we have tested the ability of PTM-Shepherd to uncover cysteine artefacts in proteomic datasets. Bekker-Jensen et al. (28) rigorously tested shotgun proteomics protocols to determine an optimal strategy for rapidly generating a comprehensive profile of human proteomes, ultimately producing a valuable repository of high-quality, deep proteomics data. Their protocol also included a 10 mM treatment with the alkylating agent CAA, which, per Schnatbaum et al. (27), only achieved two-thirds the alkylating efficiency of 10 mM IAA in complex mixtures. Unlike the other samples we analyzed for this manuscript, this protocol also did not denature protein samples before adding the alkylating agent. This likely contributes heavily to underalkylation as well. As such, it presents an exceptional opportunity to examine these cysteine artefacts.

Open search analysis followed by PTM-Shepherd shows a number of prevalent mass shifts enriched on Cys, consistent with what we expect from underalkylated samples. Note that because Cys alkylation was searched as a fixed modification, in order to elucidate the identity of modifications occurring on unalkylated Cys residues the mass shift must be decomposed into two components: a failed alkylation event (Δm = -57.0215 Da from the theoretical mass of the identified peptide) and the modification itself. Consider the mass shift -9.03680 Da detected in this dataset. PTM-Shepherd decomposes this mass shift into a failed alkylation event (−57 Da) and a triple oxidation of Cys to cysteic acid (Δm = +47.9847 Da). This becomes particularly important when trying to directly assess the number of failed alkylations in a sample. Strictly counting the number of -57 Da mass shifts will severely under count their total occurrences because it ignores cases where it is found in conjunction with another modification, which are very likely given Cys’s reactivity. We implemented an additional parameter in PTM-Shepherd to account for this that prioritizes user-defined modifications and allows them to identify mass shifts that do not directly correspond to entries in Unimod (13). On its own, failed alkylation was the sixth most abundant mass shift and was more prevalent than other common events that are often accounted for in closed searches, such as pyroglutamate formation, and accounted for 3.9% (3450 PSMs) of the 89281 total Cys-containing PSMs (**Supplementary Table 2A**). However, because it frequently occurs with other mass shifts as demonstrated above, we also pooled the instances in which it was annotated as one of two mass shifts on a peptide. Remarkably, the total number of failed alkylation events jumps to 20.5% (18343 PSMs) when considering all instances of failed alkylation annotations (**Figure 2c**). When considering decomposed mass shifts, failed alkylation is nearly twice as common as deamidation and four times as common as non-Met oxidation events (**Figure 2b, Supplementary Table 2B**).

After applying an abundance cutoff of 0.01% of total spectra, we detected 10 mass shifts exhibiting strong Cys localization (>10-fold enrichment) that were also annotated with a failed alkylation of Cys. These were a large portion of the broadly occurring Cys-enriched PTMs in the samples (**Supplementary Table 2A**). The most abundant of these modifications correlate with what would be expected in a poorly alkylated sample. The first and second isotopic error peaks in conjunction with unalkylated Cys were particularly abundant, accounting for 1601 combined spectra. The aforementioned -9.03680 Da - triple oxidation on Cys without alkylation - was most prevalent aside from these. Its heavily enriched localization to Cys (48.5-fold) lends credence to this compound identification that would be missed by other annotation tools and, consequently, a count of total failed alkylation events. Surprisingly, failed alkylation combined with a “formaldehyde adduct” (+12 Da) was also common. The combined mass shift of -45 Da was heavily localized to Cys and had a 96% N-terminal localization rate, pointing to potential thiazoladine formation via N-terminal Cys cyclization. These are known to occur in formaldehyde treated data, however the authors did not report the use of formaldehyde (29). A lone “formaldehyde adduct” mass shift accounting for 305 PSMs and heavily localized to Trp (42.5-fold enrichment) was also detected in the dataset, however. Taken together, these indicate that the thiazoladine was probably an artifact of formaldehyde exposure rather than underalkylation, though the latter may be required for there to be available Cys to react with formaldehyde. Failed alkylation of Cys and the subsequent triple and double oxidations conform to our chemical knowledge of Cys artefacts and, along with glutathione disulfide as a biological modification, comprise 8.7% of all Cys-containing PSMs. Including these modifications should increase Cys-peptide recovery in underalkylated samples.

### PTM-Shepherd computed metrics facilitate granular PTM identification

Open searches are inherently limited in the information they provide, providing only peptide lists and their associated mass shifts (4). Data interpretation efforts are further complicated by the ambiguity of mass shifts. Two methylation events and an ethylation event, for instance, would be indistinguishable from each other based on mass. However, more granular identities can be discerned by incorporating additional metrics: changes in retention time (RT), spectral similarity (SS), and localization. To demonstrate that these additional metrics improve open search result comprehension, we analyzed the synthetic unmodified tryptic peptide dataset generated as part of the ProteomeTools project (17). This dataset allows us to examine and characterize instrumental artefacts apart from confounding biological factors.

Neutral losses from peptides result from low-energy fragmentation pathways that can occur during tandem MS as well as during ionization and transmission, resulting in artefactual changes to the observed precursor mass (30). Because neutral losses occur after column elution and consequent retention time recording, they have no effect on peptide retention time. This property can be used to distinguish them from sample modifications (31). We used PTM-Shepherd to elucidate the origins of two of this dataset’s most common mass shifts attributable to both neutral losses and real modifications: loss of H_2_O and loss of NH_3_ (**Supplementary Table 3**). Peptides with multiple spectra corresponding to each loss had their RT shifts pooled and collapsed to their median.

Interestingly, both losses of H_2_O and NH_3_ exhibited bimodal changes in retention time (**Figure 3**). For peptides presenting losses of H_2_O (Δm = -18.0104 Da; 2961 peptides), both composite RT distributions were approximately Gaussian with approximate means of 0 and 450 s. As anticipated, many peptides (16.7%) fall within the mean 0 distribution, indicating that they do not experience increases in column RT despite the loss of a highly polar group. This is characteristic of neutral losses and indicates that peptides within this distribution are exhibiting neutral loss of H_2_O in the mass spectrometer prior to precursor selection. Peptides presenting losses of NH_3_ (Δm = -17.0270 Da, n = 1094) showed a similar pattern. Note that H_2_O and NH_3_ are only two examples of neutral loss and, in some cases, entire residues can be lost via this mechanism. As a significant source of instrumental bias, it is important to be able to classify neutral losses properly and remove them from experimental sample pools. In fact, for researchers studying the isobaric biological forms of these mass shifts, it’s critical to exclude these.

**Figure 3:**
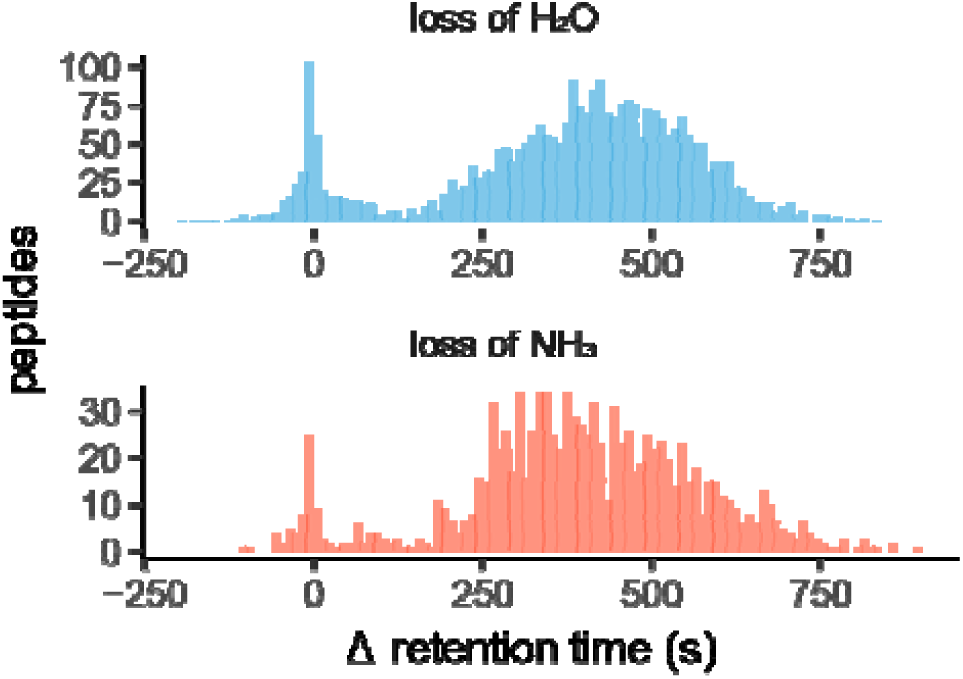
Retention time profiles for peptides with losses of H_2_O and NH_3_. Modified peptides a compared to their homologous unmodified peptides, with multiple retention time changes being collapsed to their median. The effect on retention time for losses of H_2_O (top) and NH_3_ (bottom) are distributed bimodally. These mass shifts are known to correspond to both in-source losses and spontaneous conversions. In-source losses should not have an effect on retention time, and as such are suspected to fall within a Gaussian distribution centered at zero.

The second population of peptides with H_2_O and NH_3_ loss exhibited a RT-shift consistent with a pre-elution modification - the loss of a polar group increased retention time by an average of 450 s. H_2_O losses are known to manifest as a conversion to pyroglutamic acid from Glu (32) as well as on Asn, Gln, Ser, Thr, Tyr, Asp, and Cys as sample-derived modifications (13). NH_3_ losses are known to manifest as a conversion to pyroglutamic acid, but from Gln rather than Glu (33). Other losses of NH_3_ are known to occur on some N-termini occupied by Thr, Ser, and Cys and on any Asn (13). While RT shifts alone do not contain enough information to fully identify these modifications, additional metrics calculated by PTM-Shepherd - localization propensity and modified-to-unmodified peptide similarity - allowed us to delineate the primary sources of this mass shifts.

In the case of a loss of NH_3_, two primary sources (in addition to the neutral losses) were identified: a spontaneous conversion of Gln to pyro-glutamine, and a cyclization of Cys. Cys cyclization is expected to be present in peptides being synthesized with carbamidomethylated Cys, as the reaction is known to occur after alkylation (34). Localization analysis showed that Cys (enrichment score = 10.4) and Gln (enrichment score = 4.7) were the two most enriched residues for this mass shift (**Figure 4a**). Neutral losses localized to Cys are rare and neutral losses localized to Gln are relatively common (35), which is reflected in the number of peptides each residues produces with ΔRT = 0. To illustrate, Cys has an RT profile very different from other residues in aggregate, while Gln has a similar distribution (**Figure 4c**); none of those containing Cys localized NH_3_ losses were neutral losses, as compared to 19.6% of other residues in aggregate.

**Figure 4:**
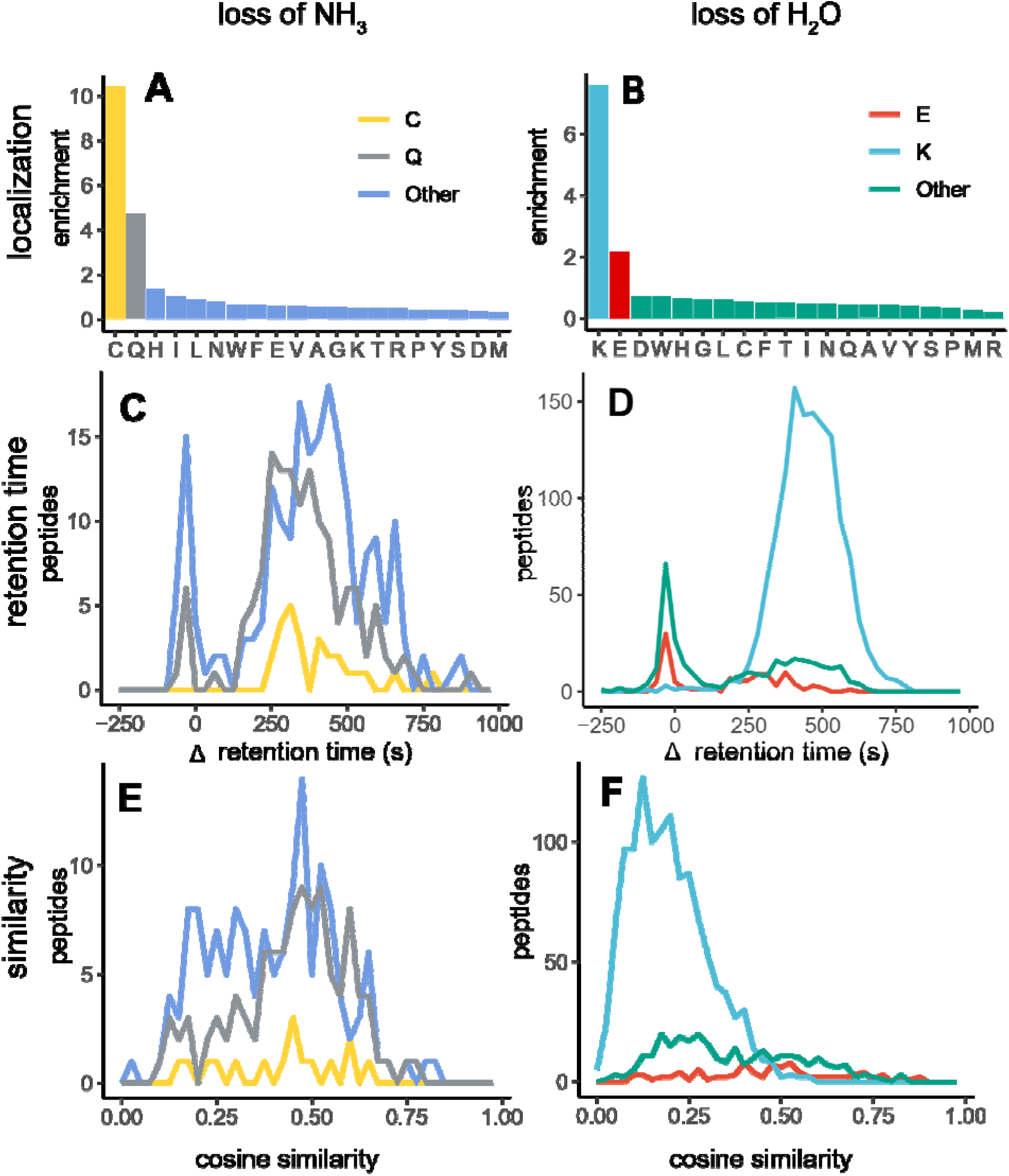
Analytical profiles for losses of H_2_O and NH_3_. A,B: Localization profiles reveal a non homogeneous landscape with specific residues showing enrichment. C,D: Select modifications are distinguishable from background in-source decays in their effect on retention time. E,F: Similarity scores show lower profiles for C-terminal modifications on Lys, whereas N-terminal modifications on Glu, Cys, and Gln have higher similarity.

Unlike NH_3_, we expected losses of H_2_O to only be heavily enriched in glutamate. Glutamate is known to have two sources for loss of H_2_O; it is known to be a source of water neutral loss but can also spontaneously undergo N-terminal cyclization *in vitro* to produce the same mass shift (36). The localization enrichment profile for loss of H_2_O (**Figure 4b**), however, revealed that two residues were exceptional contributors to this PTM’s prevalence: Glu (enrichment score = 2.2) and Lys (enrichment score = 7.6). We identified populations of peptides with losses of H_2_O corresponding to both of these populations based on ΔRT as described above (**Figure 4b**, red). An intense peak near ΔRT = 0s indicates that many unique peptides are capable of producing neutral losses of H_2_O (22.8%) on Glu, consistent with the conclusions of Sun et al. (35) Another peak near ΔRT = 300s indicates that the remainder of the peptides had a loss of H_2_O that was present before retention time was calculated, and as such is likely to be an N-terminal cyclization of glutamate occurring in vitro.

Even more so than glutamate, lysine was the largest contributor to losses of H_2_O (example spectrum at **Supplemental Figure 2**). Puzzlingly, lysine’s side chain does not have a hydroxyl group that it can readily lose, and as such any H_2_O losses attributable to lysine must be derived from the C-terminal hydroxyl group of tryptic peptides. It is worth noting that this phenomenon is unique to lysine, as Arg had the lowest localization enrichment score (0.2) of all 20 residues (**Figure 4b**). Based on retention times, there were also no appreciable H_2_O neutral losses attributable to lysine - consistent with previous findings (35) - indicating that these lysines were being dehydrated prior to retention time calculation (**Figure 4d**). We believe this is most likely due to a C-terminal lysine cyclization event. Though undescribed in proteomics, lysine derivative cyclization has been induced in other settings(37). This theory is supported by spectral similarity calculations between peptides with and without this lysine-localized mass shift (**Figure 4f**). There is exceptionally low spectral similarity between the spectra of modified and unmodified peptides. Peptides containing non-labile modifications and modifications near the C-terminus both result in low spectral similarity; non-labile modifications are likely to be retained in the MS/MS spectra rather than being removed during MS1 analysis, and C-terminal modifications shift the intense y-ion series. A covalently bound lysine ring structure on peptide C-termini fits both of these criteria and may be the underlying cause of low spectral similarity.

The similarity profiles for losses on Glu, Cys, and Gln were distinct from Lys in that they were not enriched for peptides with MS/MS showing low similarity to their unmodified counterparts.

This may be accounted for by the fact that Glu cyclization, like Cys and Gln, occurs at the N-terminus; shifting the b-ion series has less of an effect on the MS/MS spectra than shifting the y-ion series. Unsurprisingly, all three of these have similarity profiles roughly corresponding to the proportion of their spectra experiencing *in vitro* modifications, which are the only modifications we can be sure are occurring at the N-terminus.

Overall, by including metrics beyond mass shifts in PTM identification, we show that much more information beyond the chemical composition of a mass shift can be deduced. Retention times can be used to discriminate between neutral losses and sample modifications, localization profiles can be used to deduce biological or chemical origins, and spectral similarity provides additional localization and lability metrics. Incorporating all of these, we found, to our knowledge, a previously unknown or at least underappreciated modification (C-terminal lysine cyclization) in a deeply consequential synthetic peptide library.

### PTM-Shepherd in multi-experiment settings

PTM-Shepherd can be run in a multi-experiment mode to analyze modification profile across a large number of experiments. Such an analysis could be performed for visualization of interesting biological trends and in search of experiment-specific biological modifications. It could also be useful for quality control and detection of batch effects – a common source of variation in high throughput data (38). Previous efforts have been made to identify MS performance metrics (39), and some groups have shown how these can be leveraged to identify quality control issues (40) and better understand intra- and inter-laboratory variability (41, 42). We posited that open-search derived modification profiles could be used to determine interexperiment variation while simultaneously providing insight into its origins.

To evaluate PTM profiling in multi-experiment settings we used CPTAC CompRef reference material data (pooled tumor xenografts comprising ten samples each from two different breast cancer subtypes, cryopulverized and shipped to different processing locations) obtained from the CPTAC Data Portal (see **Methods**). The samples were processed at three different locations (PNNL, JHU, BI), and analyzed using TMT 10-plex labeling technology as part of the CPTAC3 Harmonization study (19). The same CompRef samples were also analyzed as longitudinal QC samples as part of the three large cancer profiling studies, CCRCC (43) (MS data collected at JHU; UCEC (44) (MS data collected at PNNL), and LUAD (MS data collected at BI).

We first investigated the most abundant mass shifts identified by MSFragger and PTM-Shepherd in these data (**Table 1**, see **Supplementary Table 4A** for the full list), which revealed several interesting observations. In general, PTM-Shepherd accurately reconstructed expected trends including the localization profiles of the most abundant modifications. For example, carbamylation and formylation were most highly enriched on N-terminus, phosphorylation on Ser, and oxidation on Trp. Not considering isotope errors, the mass shift of 229.163 (TMT overlabeling) was the most common modification, localized predominantly to Ser (enrichment factor of 5.6). Of note, only 74.5% percent of peptides found with TMT on Ser were also found in “unmodified” form (i.e., with unlabeled Ser). In contrast, many other abundant modifications, such as formylation and carbamylation, were found in both modified and unmodified forms in almost all cases. Interestingly, the second most abundant modification was a mass shift of 15.01, predominantly localized to Met (that was indistinguishable in MSFragger output from a combination of oxidation and -0.984 loss on Met). This mass shift may represent the addition of an NH group to Met due to exposure to hydroxylamine, a reagent used in TMT labeling. At present, UniMod database annotates 15.01 mass shift only as conversion of carboxylic acid to hydroxamic acid, with Asp and Glu as only possible sites (which were observed in these data, but at a lower frequency than on Met, see **Table 1**).

The PTM profiles resulting from PTM-Shepherd analysis of these data are presented in **Figure 5**. Sample-wise K-means clustering revealed distinct sample clusters, and mass shift-wise clustering on correlation between rows revealed some highly similar modifications. Sample clustering precisely reconstitutes sample processing location. For example, Cluster 2 in **Figure 5** shows a series of mass shifts related to TMT overlabeling, or TMT labeling that was not captured by fixed sequence expansion on Lys and dynamic sequence expansion on peptide N-termini. PNNL data consistently shows lower TMT overlabeling than BI and JHU for every mass shift in this cluster, and PSMs corresponding to a single additional TMT are 5-8 times lower than at the other two locations. BI and JHU also show enrichments of TMT labeling on Ser and, to lesser degree, Thr.

**Figure 5:**
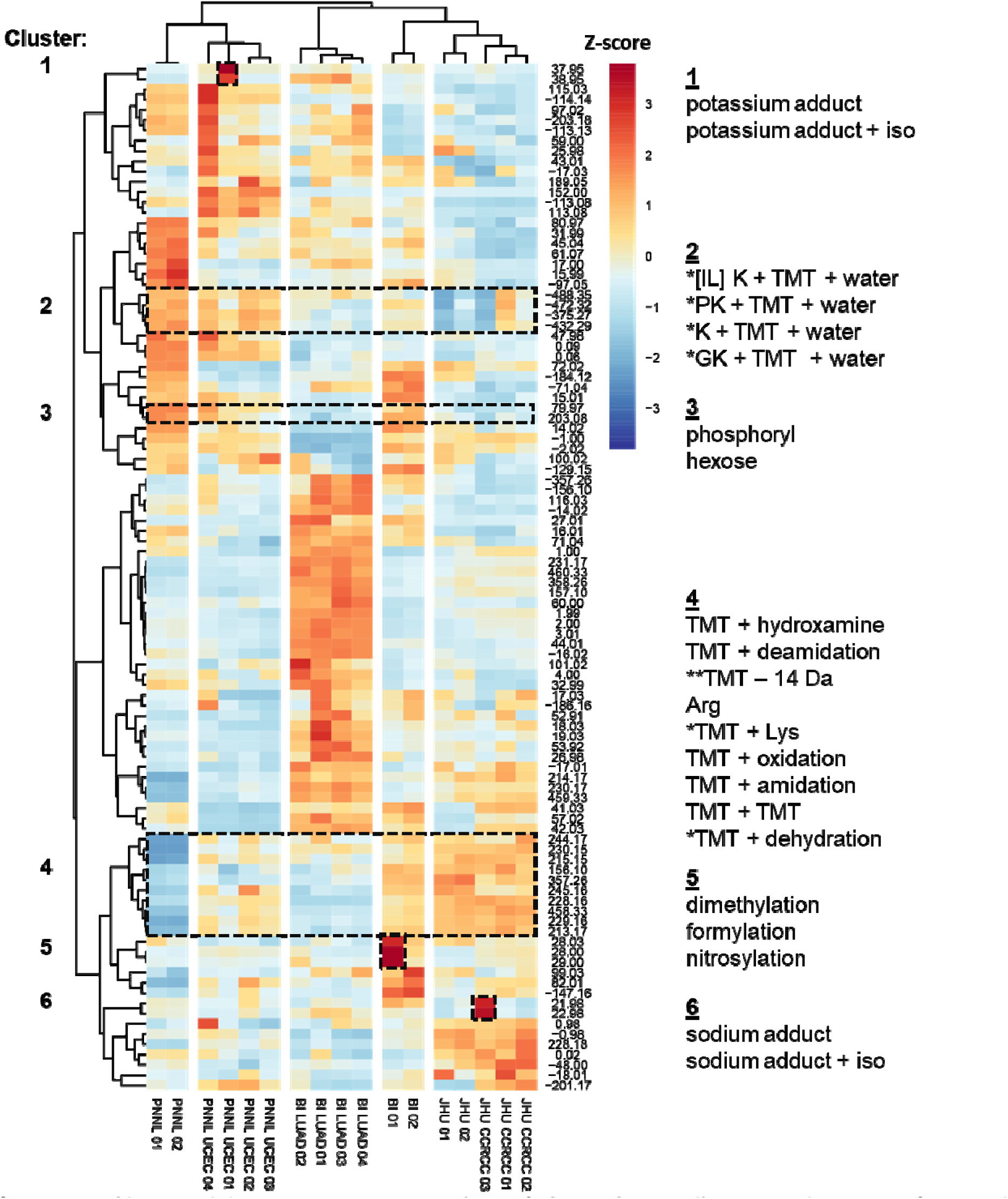
Clustered heatmap representation of CPTAC3 quality control xenograft samples. Values shown are row-wise z-scores of spectral counts. Row clustering shows highly related modifications, and column clustering shows experiments clustering by processing location. Mass shift clusters are numbered and their corresponding mass shifts are shown top-to-bottom in the to the right of the figure. Samples with no tumor type label (LUAD, UCEC, CCRCC) were processed as part of the CPTAC harmonization study. Other samples were processed longitudinally throughout their respective studies. *This annotation was constructed manually. **-14 Da can correspond to a large number of modifications and single residue mutations.

Though we expect to see differences in TMT labeling fidelity, PTM-Shepherd also allows us to explore unexpected batch effects. Lenčo *et al*. recently raised concerns about the use of formic acid in sample preparation, specifically stating that an excess of Ser, Thr, and N-terminal formylation events are present in samples reconstituted with it (45). Within the CPTAC harmonization study, the localization profile did exactly match that described by Lenčo and colleagues: Ser enrichment of 2.7, Thr enrichment of 1.9, and a 61% potential N-terminal rate (**Supplementary Table 4A)**. Interestingly, this formylation peak appears to disproportionately high in the first of the two BI replicates from the harmonization studies (**Figure 5**, Cluster 5). BI01 replicate had formylation 10- to 20-fold higher than JHU01 or PNNL01 and was even 4-fold higher than BI02. Ser and Thr also exhibit inflated formylation localization for this sample, consistent with Lenčo *et al*.’s results (**Supplementary Table 4B-C)**. Overall, a deeper analysis may be warranted in future studies to reduce batch effects caused by formic acid use. These analyses could be extended to other sample handling artifacts, e.g. potassium adducts (Cluster 1) and sodium adducts (Cluster 6), which also exhibit marked longitudinal variability.

PTM-Shepherd also reveals how instrument parameters might be playing a role in lab-specific batch effects. We noted four apparent C-terminal neutral losses that were differentially identified across labs (**Figure 5**, Cluster 2). Sequence analysis facilitated manual decomposition, revealing that three of the mass shifts were composed of a hydrophobic residue (Iso, Leu, Pro, or Gly), a C-terminal Lys, a TMT tag, and a water. The final mass shift was the loss of a C-terminal Lys, a TMT tag, and a water. Interestingly, most of PSMs exhibiting these shifts also had N-terminal TMT labels, indicating that there may be a relation between C-terminal neutral losses and N-terminal TMT labeling for some peptides. It’s possible that slight differences in fragmentation or ionization energies between labs may be manifesting as disparities in precursor charge states and proton mobility, however determining the mechanism through which these are occurring is outside the scope of this work.

Aside from analyzing how samples cluster together, there is also useful information to be gleaned from analyzing how mass shifts cluster together. We expect that related modifications should have highly correlated abundances between experiments, with co-clustering of isotope peaks as the most obvious example (e.g., potassium and sodium adducts and their related first isotopic peaks, **Figure 5**, Clusters 1 and 6). Using a *Z*-score normalization to coerce PTMs derived from related sources across experiments to cluster together and clustering on the correlation between rows, we were able to identify some unknown mass shifts based on their co-clustering with other (known) modifications. In the TMT-related cluster noted above (**Figure 5**, Cluster 2), we observed three mass shifts that were missed by automatic annotation. Of the six that were annotated, five of the mass shifts are directly attributable to TMT overlabeling, and one to a missed tryptic cleavage or additional Arg at one end of the peptide sequence. This knowledge allowed us to explain one of the unannotated mass shifts as combinations of missed cleavages and TMT labeling. One modification (+357.2584 Da) is precisely the mass of a TMT-labeled Lys residue. This was missed by automatic annotation because both “Addition of Lys” and “TMT10 Plex” would have to be more abundant than the combination of the two during PTM-Shepherd analysis. The other unexplained mass shift (+213.1680 Da) can be explained by a combination of TMT overlabeling and a dehydration event or a misattributed Met oxidation included as a variable modification. Overall, this analysis demonstrates the utility of multi-experimental analysis with PTM-Shepherd to better identify the mass shifts are not amenable to automatic annotation.

Finally, while our analysis above focused mostly on modifications introduced due to labeling and other sample handling steps, clustering of mass shifts may also be useful for uncovering correlated biological modifications. Of note, clustering of PTM-Shepherd results shows that phosphorylation (+79.97 Da) is most correlated with HexNAc (+203.08 Da) (**Figure 5**, Cluster 3). Interestingly, co-enrichment of glycopeptides was recently observed in datasets experimentally enriched for phosphopeptides (46), however, no phosphopeptide enrichment steps were applied to generate the data used in this work.

## CONCLUSIONS

Despite advancements in computational proteomics, many MS/MS spectra remain unexplained. Open searching with tools like MSFragger has proven to be an effective way to overcome the limitations of traditional database searches by removing the requirement of having prior knowledge of the peptide modifications present in the sample. The modifications elucidated by open searches, however, lack many of the metrics necessary to make proper determinations about their identities and origins. We addressed these challenges in PTM-Shepherd, which produces comprehensive PTM profiles for open search derived mass shifts, including multiple Unimod annotations, retention time changes, spectral similarity, and localization profiles.

We demonstrated the utility of PTM-Shepherd in four examples, providing a broadly applicable guide for others interested in utilizing open searches for PTM analysis in their own research. First, in the development of an FFPE-treatment PTM palette, we showed how PTM-Shepherd disambiguated two overlapping peaks: formylation and demethylation. We also demonstrated how PTM palettes can be easily constructed for other sample preparation methods without extensive postprocessing. Second, we showed how PTM-Shepherd’s unique ability to decompose mass shifts into multiple Unimod modifications allows us to identify and quantify the degree of failed alkylation, though this is easily extensible to other scenarios, e.g., identifying mass shifts corresponding to an absent variable modification and another co-occurring modification. We also demonstrated how incorporating additional metrics into PTM identification provides researchers with more granular, high confidence PTM identities, including the ability to distinguish between sample-derived and instrument-derived artefacts. Finally, when applied to data from a large, multi-center proteomics study, PTM-Shepherd helped us to visualize batch effects and the effect of sample processing location, as well as elucidate the identities of unannotated mass shifts. We believe PTM-Shepherd will become a widely used component in our MSFragger-based pipeline for comprehensive analysis of post-translational and chemical modifications, including searches for rare and even novel modifications, across a wide range of biological applications.

## Acknowledgements

The authors would like to thank the users of our tools for their feedback. This work was funded in part by NIH grants R01-GM-094231 and U24-CA210967. D.J.G. was supported in part by the Advanced Proteome Informatics of Cancer training program (T32 CA140044).

## Data and Software Availability

All raw mass spectrometry data used in the manuscript can be found from the ProteomeXchange Consortium via the PRIDE partner repository, or (CPTAC data) from the CPTAC Data Portal, using specific dataset identifiers cited in the text. PTM-Shepherd is available as a standalone JAR executable (https://github.com/Nesvilab/PTM-Shepherd) and also fully integrated in FragPipe Graphical User Interface (http://fragpipe.nesvilab.org/).

## Author Contributions

A.I.N. and A.T.K. conceived the project, A.T.K. developed the first version of the software, later extended and improved by D.J.G., D.J.G. performed all analyses, D.M.A., F.Y., H.Y.C., and F.L. contributed to the software development, D.J.G and A.I.N. wrote the manuscript with input from all authors, and A.I.N. supervised the entire project.

## Abbreviations

PTM: post-translational modification
PSM: peptide-spectrum-match
FFPE: formaldehyde-fixed paraffin-embedded
FDR: false discovery rate
CPTAC: Clinical Proteomics Tumor Analysis Consortium
TMT: tandem mass tag
BI: [the] Broad Institute
JHU: Johns Hopkins University
PNNL: Pacific Northwest National Lab
CCRCC: clear cell renal cell carcinoma
GBM: glioblastoma
LUAD: lung adenocarcinoma
UCEC: uterine corpus endometrial carcinoma
CAA: chloroacetamide
IAA: iodoacetamide
RT: retention time

