## Supplemental Figure 1, Supplemental Figure 2, and table legends for "PTM-Shepherd: analysis and summarization of post-translational and chemical modifications from open search results"

Supplementary Materials

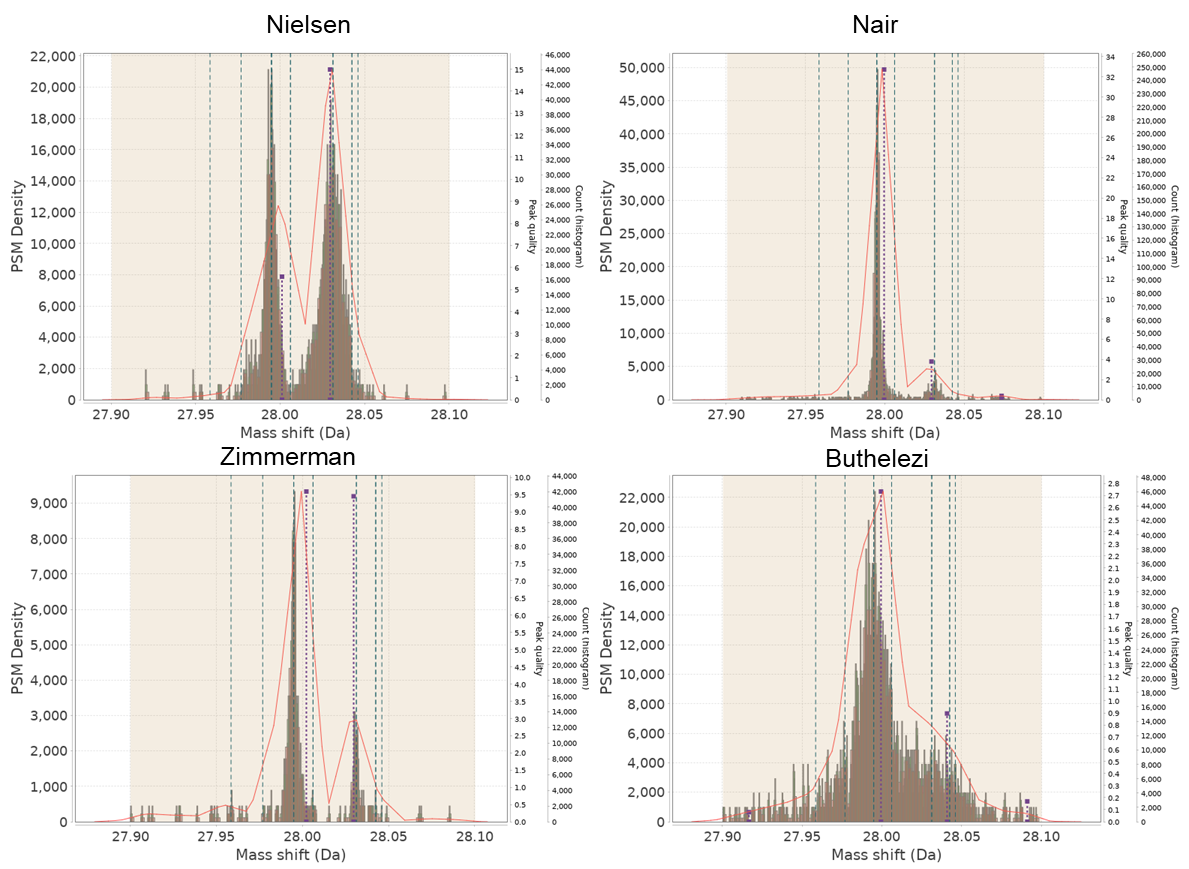

**Figure S1:** DeltaMass mass shift profiles for the 27.90 to 28.10 Da region for FFPE treated data. Two peaks representing formylation (+27.99 Da) and di-methylation (+28.03 Da) are clearly visible for the Nielsen, Nair, and Zimmerman datasets. For the lower resolution Buthelezi dataset, the composite gaussian mixture (red outline) is still composed of two distributions centered at the expected values (purple squares).

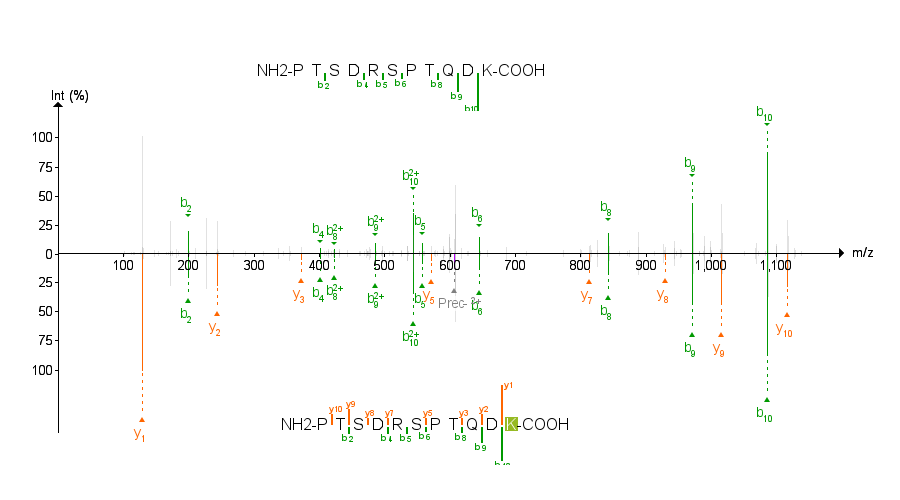

**Figure S2:** PDV view of spectrum 01717a_BE3-TUM_second_pool33_01_01-3xHCD-1h-R1.6388.6388.2. Placing the loss of water (-18.0106 Da) on C-terminal K aligns the *y*-ion series. All discriminating ions to confirm localization are visible.

Supplementary Tables 1-4 contain lightly formatted output from PTM-Shepherd. Four table tables are included, with their column annotations as follows:

- ${DATASET}.profile.tsv: important summary statistics of the dataset (or all datasets combined)
  - Peak Apex: apex of the detected peak (Da)
  - Peak Lower: lower bound of the detected peak (Da), determined by precursor tolerance or the detection of an adjacent peak
  - Peak Upper: upper bound of the detected peak (Da), determined by precursor tolerance or the detection of an adjacent peak
  - PSMs:	the number of PSMs contained within the peak boundary. This is reported for each dataset if multiple datasets are used as input
  - PSMs/million: the number of PSMs over the number of scans collected in the experiment. This is reported for each dataset if multiple datasets are used as input.
  - Peptides: the number of unique peptides containing the mass shift
  - % in unmodified: the percentage of PSMs in this mass bin with a corresponding PSM in the unmodified bin. This is reported for each dataset if multiple datasets are used as input.
  - Potential Modification 1 / 2: modification annotations derived from Unimod. Isobaric modifications are denoted with a “/” between them. The combination of the two modifications adds up to the total mass shift
  - Similarity (mean): modified peptides are compared to their unmodified counterparts. When multiple modified-unmodified comparisons are done for a single peptide, the cosine similarity similarity scores are averaged for the peptide. The peptide scores are then averaged across all peptides in the mass shift bin. These comparisons are only done for peptides in the same charge state.
  - Similarity (variance): variance in peptide similarity scores discusses above
  - DeltaRT (mean): modified peptides are compared to their unmodified counterparts. When multiple modified-unmodified comparisons are done for a single peptide, the changes in retention times are averaged for the peptide. The peptide scores are then averaged across all peptides in the mass shift bin. These are only done for peptides in the same run.
  - DeltaRT (variance):	variance in peptide changes in retention time scores discussed above
  - Localized PSMs: number of PSMs in the mass bin that showed at least one additional matched ion when the mass shift is placed on a residue
  - N-term rate: percentage of PSMs with an uninterrupted string of localized residues from the N-terminus. This is calculated differently from other enrichment scores due to the difference in assumptions underlying N-terminal and residue-specific localization.
  - Residue enrichment scores:	The first number reported is the enrichment score, roughly equivalent to the odds of being localized to this residue vs other residues. The second number reported is the weighted number of PSMs where the mass shift was localized to this residue. Shifts localizing to multiple residues are divided by the number of localized residues in the spectra.
- global.modsummary.tsv: summary of estimated modification counts after decomposition into multiple modifications
  - Modification: name matched to mass shift during annotation
  - Theoretical mass shift: Theoretical mass shoft produced by modification. Unannotated mass shifts show their experimental mass shifts instead of their theoretical ones
  - PSMs/million: Summed PSMs/million for every mass shift where the modification was identified as either the single mass shift or one of the two decomposed mass shifts
  - PSMs: Summed PSMs for every mass shift where the modification was identified as either the single mass shift or one of the two decomposed mass shifts

**Supplementary Table 1:** PTM-Shepherd results for reanalysis of four Tabb et al. (2020) datasets. See “Supplementary_table_S1.xlsx”.

- global.profile.tsv table from PTM-Shepherd containing summary statistics across all four datasets

**Supplementary Table 2:** PTM-Shepherd results for reanalysis of Bekker-Jensen et al. (2017) dataset. See “Supplementary_table_S2.xlsx”.

- **A:** global.profile.tsv table from PTM-Shepherd containing summary statistics across the dataset
- **B:** global.modsummary.tsv table from PTM-Shepherd containing abundance information for annotated modifications rather than mass shifts

**Supplementary Table 3:** PTM-Shepherd results for reanalysis of Zolg et al. (2017) dataset. See “Supplementary_table_S3.xlsx”.

- global.profile.tsv table from PTM-Shepherd containing summary statistics across the dataset

**Supplementary Table 4:** PTM-Shepherd results for reanalysis of CPTAC3 quality control samples. See “Supplementary_table_S4.xlsx”.

- **A:** global.profile.tsv table from PTM-Shepherd containing summary statistics across the datasets
- **B:** 01BI.profile.tsv report from PTM-Shepherd containing summary statistics for this dataset
- **C:** 02BI.profile.tsv report from PTM-Shepherd containing summary statistics for this dataset
